# Proteomic profiling of tissue extracellular vesicles (EVs) identifies tissue-specific EV markers and predicts the accessibility of tissue EVs to the circulation

**DOI:** 10.1101/2025.03.06.641016

**Authors:** Sho Watanabe, Ryuichiro Sato, Takashi Sasaki, Yu Takahashi, Yoshio Yamauchi

**Affiliations:** Laboratory of Food Biochemistry, Department of Applied Biological Chemistry, Graduate School of Agricultural and Life Sciences, The University of Tokyo, Tokyo, Japan; Nutri-Life Science Laboratory, Department of Applied Biological Chemistry, Graduate School of Agricultural and Life Sciences, The University of Tokyo, Tokyo, Japan

**Keywords:** extracellular vesicles (EVs), exosomes, mouse tissues, proteomics

## Abstract

Extracellular vesicles (EVs) mediate cell-to-cell communication in endocrine, paracrine, and autocrine fashions, but their roles and transport in the body remain incompletely understood. It is expected that EVs circulate in the body through the bloodstream, where EVs derived from various tissues are present. However, how much each tissue contributes to circulating EVs is largely unknown. Therefore, identifying tissue-specific EV markers is of importance for elucidating *in vivo* dynamics of tissue EVs. A key issue that hampers studying EVs *in vivo* is the lack of methodologies for collecting them from tissues. Here, we developed methods to isolate EVs from four different tissues, skeletal muscle, heart, liver, and adipose tissue, and performed proteomic analysis on their EVs. Unbiased and quantitative proteomic analysis revealed protein signatures of EVs from the four tissues and identified marker proteins specific to EVs of each tissue. Furthermore, through comprehensive comparisons of the proteome from tissue and plasma EVs, we predicted the accessibility of EVs from the four tissues to the circulation. Our data suggests that, among the four tissues, adipose tissue-derived EVs more readily enter the circulation, while skeletal muscle-derived EVs exhibit poor accessibility to the blood. Collectively, our tissue EV proteome identified tissue-specific EV markers and suggests that the accessibility of tissue EVs to circulation highly depends on the original tissues.

## Introduction

Extracellular vesicles (EVs) are released from all tissues and cell types and play a role in intercellular communication by transporting diverse biomolecules, including nucleic acids (dsDNA, mRNA, microRNA, and long noncoding RNA), proteins, and lipids to recipient cells [1–5]. Upon delivery to recipients, these biomolecules exert a wide range of biological functions [6, 7]. EVs are categorized into three classes based on their size and biogenesis mechanisms: exosomes (50–150 nm in diameter), microvesicles (300–1,000 nm), and apoptotic bodies (100–5,000 nm) [1–3]. Once released from donor cells, EVs function in autocrine and paracrine fashions within the original tissue and are also transported to remote tissues through systemic circulation, modulating physiology and pathophysiology of recipient tissues and cells [8–10]. In addition, due to their stability and presence in various body fluids such as blood and urine, EVs are utilized for therapeutic and diagnostic purposes [11, 12].

Determining the contents of EVs is crucial for understanding their roles. Unbiased omics analyses have been employed to uncover the molecular signatures of EVs released from various cell types [13–15]. In particular, proteomic analysis has been performed to identify universal and cell type-specific EV marker proteins [16–19]. Such marker proteins may serve as tools for identifying and isolating specific EV subpopulations *in vivo*. We performed proteomic analysis of small EVs released from mouse and human skeletal muscle cells and identified several muscle-specific proteins that serve as potential skeletal muscle (SkM)-derived EV markers *in vivo* [19]. Other investigators also revealed proteomic profiles of small EVs derived from cell lines of various tissue origins, including adipocytes, endothelial cells, hepatocytes, and myocytes, and proposed adiponectin as an adipose tissue-derived EV marker [18]. Although cell-based studies have identified potential tissue-specific EV marker proteins, proteomic signatures of tissue EVs *in vivo* are largely unknown. Therefore, whether tissue-specific EV markers determined by cell-based experiments can directly be applied to *in vivo* studies remains to be validated.

Upon secretion from tissues, significant portions of EVs are expected to enter the blood and circulate in the body. Therefore, blood should contain EVs of various origins. Several studies have shown that tissue EVs are transported to other tissues and regulate the physiology of target tissues [20, 21]. However, what extent each tissue contributes to circulating EVs is largely unknown at present. Recent studies have shown that tissue interstitium contains significant amounts of EVs and suggested that tissue EVs are concentrated in a corresponding tissue [22–24]. In addition, other studies have identified key molecules involved in EV-mediated interorgan communication [25], cellular quality controls [26], and drug responses [27]. Therefore, isolation of EVs present in a tissue is a powerful approach to determining reliable tissue-specific EV markers and studying *in vivo* significance of EVs. On the other hand, there is no report on developing a unified protocol for isolating tissue EVs that is compatible with multiple tissues.

The aims of this work are to develop methods for the isolation of EVs from multiple tissues and to perform comprehensive proteomic comparisons among EVs from different tissues and between tissue and plasma EVs. By refining the previous protocols, we develop methods that efficiently collect tissue EVs from skeletal muscle, heart, liver, and adipose tissue. Based on proteomic profiling of tissue and plasma EVs, we propose tissue-specific EV marker proteins that are applicable for *in vivo* studies and predict the accessibility of EVs from these four tissues to the bloodstream.

## Results

### Isolation and characterization of tissue EVs

To perform proteomic characterization of tissue EVs, we first attempted to develop methods to isolate tissue EVs from four mouse tissues, skeletal muscle (gastrocnemius, soleus, tibialis anterior, and quadriceps), heart, liver, and adipose tissues (epididymal white adipose tissues). **Figure 1** shows the schematic flows of the methods. To isolate EVs from tissue interstitial spaces, tissues were gently minced and treated with collagenase and dispase for one hour at 37°C. Afterward, tissue homogenates were passed through a 100-μm strainer to remove tissue fragments, followed by the removal of debris by three-step centrifugation. For studying small EVs, we filtrated the supernatant containing EVs using a 0.20-μm filter to exclude EVs larger than 200 nm. Subsequently, tissue EVs were isolated by three different methods (**Figure 1**).

**Figure 1.**
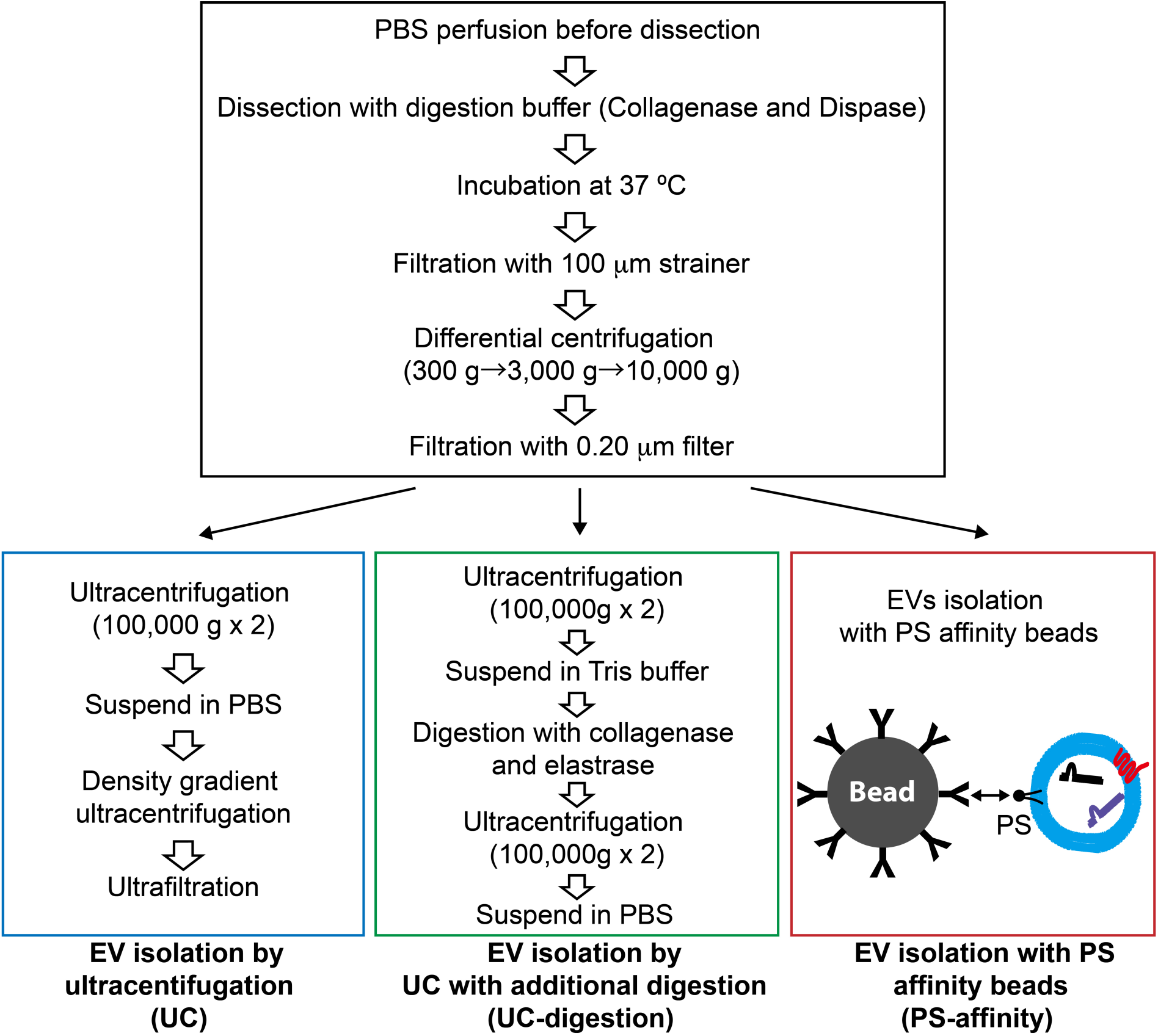
Schematic overview of three procedures for isolating tissue EVs. Tissues are sliced and treated with type II collagenase and dispase II at 37°C. After filtration and three-step centrifugation for removing debris, samples are filtrated with a 0.20-μm filter to exclude large EVs. Tissue EVs are then isolated by the three methods. In the UC method (blue frame), tissue EVs are first collected by ultracentrifugation at 100,000 g and then subjected to density-gradient ultracentrifugation with Optiprep at 200,000 g. In the UC-digestion method (green frame), tissue EVs are first collected by ultracentrifugation at 100,000 g and then treated with type II collagenase and elastase. Finally, EVs are collected by ultracentrifugation. In the PS-affinity method (red frame), tissue supernatant containing EVs is concentrated to ∼1 mL and mixed with Tim 4-Fc-bound streptavidin magnetic beads. PS^+^ EVs are pulled down and eluted.

First, we isolated skeletal muscle-derived EVs (SkM-EVs), heart-derived EVs (heart-EVs), liver-derived EVs (liver-EVs), and adipose tissue-derived EVs (adipose-EVs) by the UC method and examined the distribution of EVs by detecting the classical EV marker proteins Alix and CD81. The result showed that tissue EVs from the four tissues were recovered in fractions 4–6 with a density of 1.03–1.20 g/mL (**Figure 2A**), which is in good agreement with a previous report [28]. Transmission electron microscopy (TEM) analysis showed vesicular morphology of tissue EVs isolated from the four tissues (**Figure 2B**). The diameters of SkM-EVs, heart-EVs, liver-EVs, and adipose-EVs were 70.4 ± 31.8 nm, 66.9 ± 32.5 nm, 62.7 ± 32.2 nm, and 65.3 ± 27.6 nm, respectively, and their sizes were not different among these tissues (**Figure 2C**). We noticed that SkM-EVs contain extracellular matrix (ECM)-like fibrillar structures attached to EVs (**Figure 2B**), suggesting that the collagenase and dispase treatment in the first step was insufficient to digest ECM because skeletal muscle is an ECM-rich tissue [29, 30]. On the other hand, this observation is consistent with our previous results that EV-like vesicles are bound to ECM-like structures within this tissue [19].

**Figure 2.**
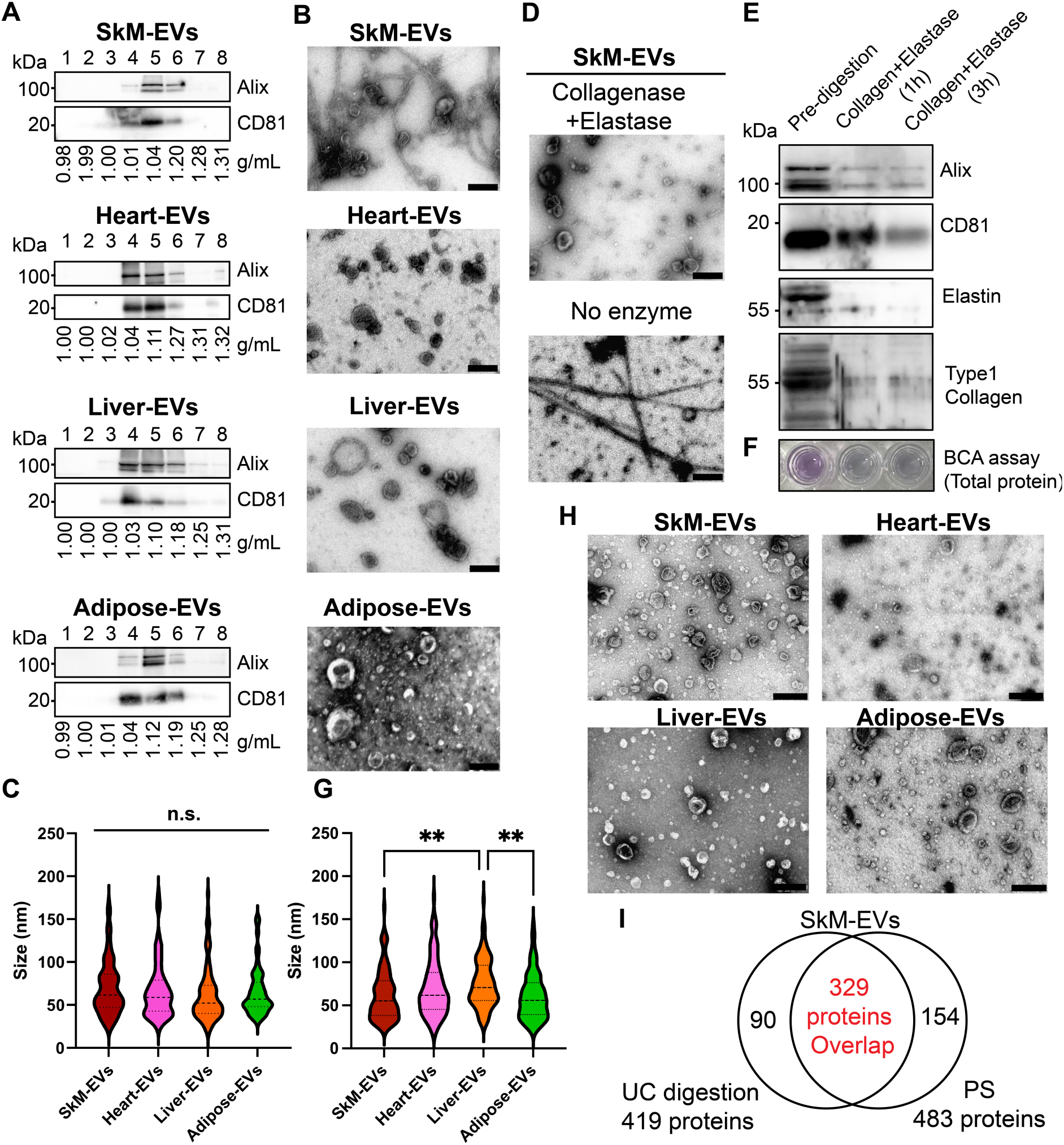
Isolation of tissue EVs from four tissues. (A) Distribution of classical EV markers. After density-gradient centrifugation of SkM, heart, liver, and adipose EVs in the UC method, aliquots of each fraction were subjected to immunoblot analysis. Alix and CD81 were used as EV marker proteins. (B) TEM images of EVs isolated from skeletal muscles, hearts, livers, and adipose tissues by the UC method. Scale bar, 200 nm. (C) Size distribution of tissue EVs isolated by the UC method. The size of EVs was measured based on TEM images. Statistical analysis was performed by one-way ANOVA with Tukey-Kramer post-hoc test. n = 100; n.s., P > 0.05. (D) TEM images of SkM-EVs isolated by the UC-digestion method. ECM-like components were largely eliminated after the treatment with collagenase and elastase. Scale bar, 200 nm. (E) Detection of ECM components before and after the additional digestion. After SkM-EVs were treated with collagenase and elastase for 1 or 3 h, SkM-EVs were collected by ultracentrifugation, and Alix, CD81, elastin, and type I collagen were detected by immunoblotting. (F) Total protein concentrations in SkM-EVs before and after the additional digestion (1 or 3 h) were measured using a micro BCA assay. Shown are wells after colorization. (G) Size distribution of tissue EVs isolated by the PS-affinity method. Statistical analysis was performed by one-way ANOVA with Tukey-Kramer post-hoc test. n = 100. **P < 0.01, *P < 0.05. (H) TEM images of tissue EVs isolated by the PS-affinity method. Scale bar, 500 nm. (I) Venn diagram showing proteomic overlap between SkM-EVs isolated by the UC-digestion and PS-affinity methods.

Since the types of collagenases used for tissue dissociation were not sufficient to digest and remove ECM (**Figure S1**), we are concerned that the contamination of ECM-like structures may affect subsequent proteomic analysis of tissue EVs. To exclude this possibility, we treated SkM-EVs with collagenases and elastase to digest collagen and elastin that are rich in skeletal muscle after the first sedimental ultracentrifugation. This method is referred to as the UC-digestion method (**Figure 1**). After the additional digestion step, SkM-EVs were collected by ultracentrifugation. We confirmed by TEM that this additional treatment significantly reduced the contamination of ECM (**Figure 2D**) and immunoblotting (**Figure 2E**). **Figure 2E** shows that one-hour treatment is sufficient to reduce elastin and type I collagen in SkM-EV samples. Total protein amounts measured by BCA assay also decreased after the additional digestion (**Figure 2F**).

Because ultracentrifugation potentially causes the aggregation of EVs [31], we also attempted to isolate tissue EVs by an ultracentrifugation-free method. We took advantage of the high-affinity binding of Tim4 to phosphatidylserine (PS), a phospholipid abundant on the surface of EVs [32]. This method is termed the PS-affinity method (**Figure 1**). By using Tim4-coated magnetic beads (MagCapture Exosome Isolation Kit PS), we previously isolated interstitial EVs from skeletal muscle [19]. We applied this method to other tissues. We observed isolated SkM-EVs, heart-EVs, liver-EVs, and adipose-EVs under TEM (**Figure 2H**) and found that their sizes are all within 30–180 nm (**Figure 2G**), confirming good isolation of small EVs from the four tissues by this method. SkM-EVs (78.6 ± 25.5 nm) and adipose-EVs (76.7 ± 22.2 nm) were slightly but significantly smaller than liver-EVs (92.5 ± 22.7 nm). The difference in the size of tissue EVs isolated by the UC method (**Figure 2C**) and PS-affinity method (**Figure 2G**) may be explained by the principle of the two methods; the UC method isolated almost all EVs in tissue interstitium while the PS-affinity method collects only PS^+^ EVs.

To further compare the biochemical nature of SkM-EVs isolated by the UC-digestion and PS-affinity methods, we performed quantitative proteomic analysis on SkM-EVs isolated by the two methods. We identified 419 proteins and 483 proteins in SkM-EVs isolated by the UC-digestion and PS-affinity methods, respectively (**Supplementary Table 1**) and found that 329 proteins (79% and 68%) overlapped between EVs isolated by the two methods (**Figure 2I**). In the proteomic data, collagens were not major proteins and elastin was undetectable, indicating that the two methods efficiently excluded ECM. We noted that in both methods, mitochondria-related proteins and -Gene Ontology (GO) terms were enriched in SkM-EVs (**Figure S2, Supplementary Table 1**), suggesting that skeletal muscle releases mitochondria-derived small vesicles as proposed [26].

### Proteomic profiling of tissue EVs

Next, we sought to determine the proteomic profile of SkM-EVs, heart-EVs, liver-EVs, and adipose-EVs (**Figure 3A**). For this purpose, we employed the PS-affinity method for avoiding ultracentrifugation that may affect the quality of EVs [31]. The quantitative shotgun proteomic analyses on SkM-EVs, heart-EVs, liver-EVs, and adipose-EVs identified 483, 578, 617, and 532 proteins, respectively, covering a total of 771 different proteins (**Figure 3B**, **Supplementary Table 2**). Of the 771 proteins, 370 proteins (48%) were found in all tissue EVs (**Figure 3B**). As mitochondrial proteins have been identified in EVs [23, 33–35], GO analyses showed that mitochondria-related proteins are enriched in all tissue EVs (**Figure 3C**), suggesting that mitochondrial vesicles are actively released from all these tissues. In addition, we found unique components that are included in a tissue-specific manner; SkM-EVs, heart-EVs, liver-EVs, and adipose-EVs contain five, 14, 71, and 16 specific proteins, respectively (**Figure 3B**). To further characterize tissue EVs, we classified their components based on the GO categories using DAVID, an integrative platform [36, 37]. Within the “Biological processes” term, proteins classified into “Muscle contraction” were enriched in SkM-EVs and heart-EVs, indicating that SkM-EVs and heart-EVs contain muscle-related proteins (**Figure 3C**). On the other hand, proteins classified into “lipid metabolic process” were found enriched in liver-EVs and adipose-EVs (**Figure 3C**). These results suggest that tissue EVs display distinct protein signatures reflecting their original tissues.

**Figure 3.**
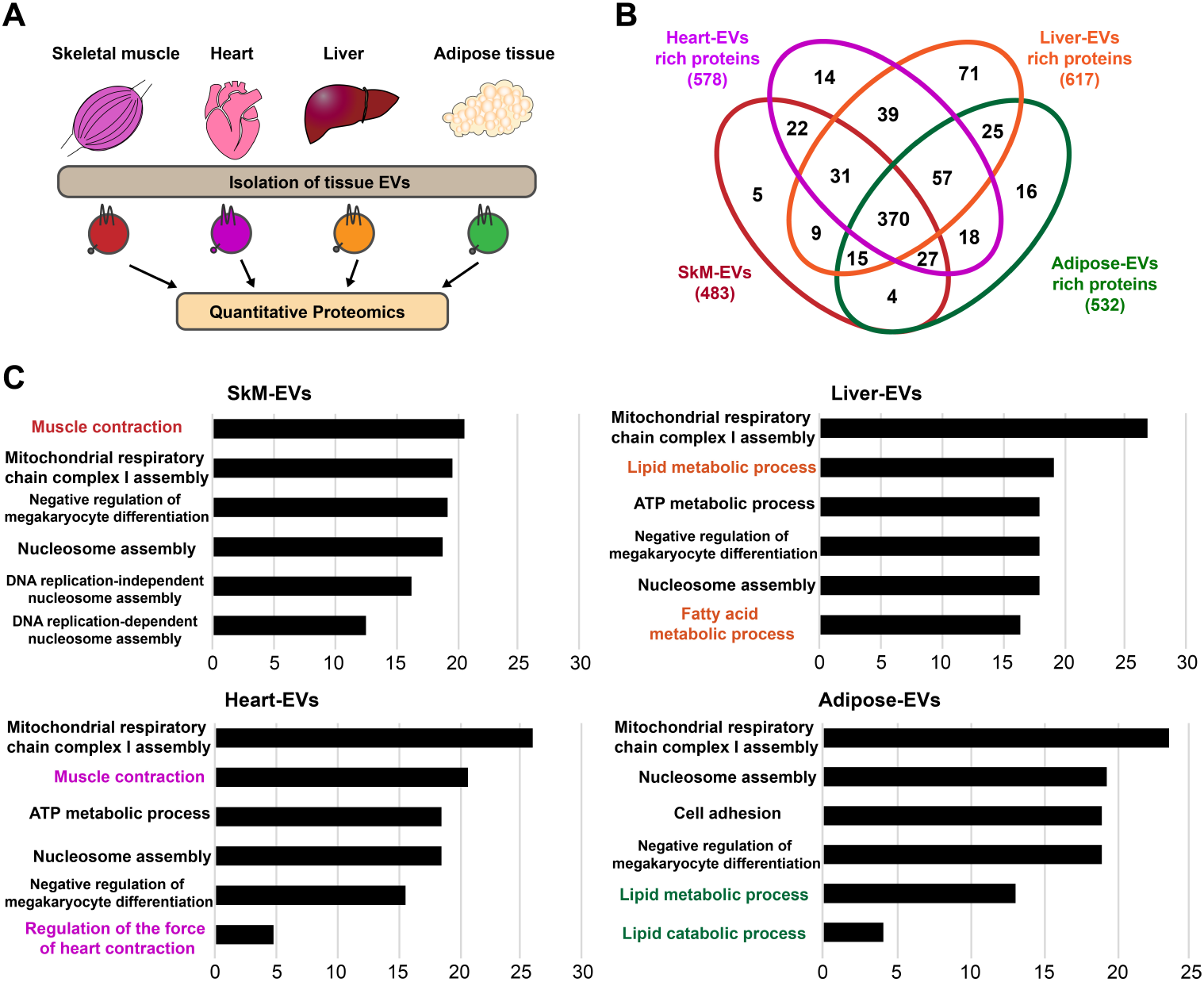
Proteomic profiling of tissue EVs isolated by the PS-affinity method. (A) A schematic experimental diagram. Tissue EVs were isolated from skeletal muscle, heart, liver, and adipose tissues by the PS-affinity method for quantitative proteomic analysis. (B) Venn diagram showing the distinct and overlapping proteins among tissue EVs isolated from the four tissues. (C) GO analysis of tissue EVs for biological processes. Proteomic data on SkM-EVs, heart-EVs, liver-EVs, and adipose-EVs were analyzed using DAVID. The top six GO terms in the enrichment analysis of biological processes are listed.

We next performed comparative analyses on our tissue EV proteomics data with proteome of EVs released from respective cultured cells. We first compared proteomes between the current SkM-EVs and C2C12 myotube-derived EVs we previously reported [19] (**Supplementary Table 3**). Of the 573 proteins in SkM-EVs isolated by the UC-digestion and PS-affinity methods, 204 proteins (approx. 36%) were found in EVs from C2C12 myotubes (**Figure S3**). The results suggest that skeletal muscle contains EVs released not only from myotubes/myofibers but also from other cell types, as this tissue is composed of various cell types, including immune cells, fibro-adipogenic cells, and endothelial cells [38]. We also compared our proteomes of SkM-EVs, liver-EVs, and adipose-EVs isolated by the PS-affinity method with published proteomics data on C2C12 myotube-derived EVs, AML12 hepatocytes-derived EVs, and 3T3L1 adipocytes-derived EVs [18], respectively. Of the 483 SkM-EV proteins, 92 proteins (19%) overlapped with EVs released from C2C12 myotubes (**Figure S4A**). Of the 617 liver-EV and 532 adipose-EV proteins, 98 (16%) and 88 proteins (16%) were found in EVs derived from AML12 hepatocytes and differentiated 3T3L1 adipocytes, respectively. The relatively low overlap between tissue EV and corresponding cell line-derived EV proteomes suggests that tissue EVs are composed of those released from various cell types.

### Identification of potential tissue-specific EV marker proteins

Tissue-specific EV marker proteins remain poorly characterized *in vivo*. To seek such EV markers, we extracted proteins enriched more than 10-fold in EVs from one tissue over those from the other three tissues. We identified 12 such proteins for SkM-EVs, 29 for heart-EVs, 136 for liver-EVs, and 42 for adipose-EVs (**Figure 4A**). We assessed their tissue specificity using a public database, The Human Protein Atlas, and found that many proteins we selected are predominantly expressed in their original tissues (7 out of 14 proteins for SkM-EVs, 10 out of 29 proteins for heart-EVs, 79 out of 136 proteins for liver-EVs, and 7 of 42 proteins for adipose-EVs) (**Supplementary Table 4**). These tissue EV-specific proteins rarely overlapped with EV proteins from respective cell lines reported previously [18] (**Figure S4B**). These results suggest that attention is needed when protein signatures of EVs determined by cell-based studies are compared to those of tissue EVs *in vivo*.

**Figure 4.**
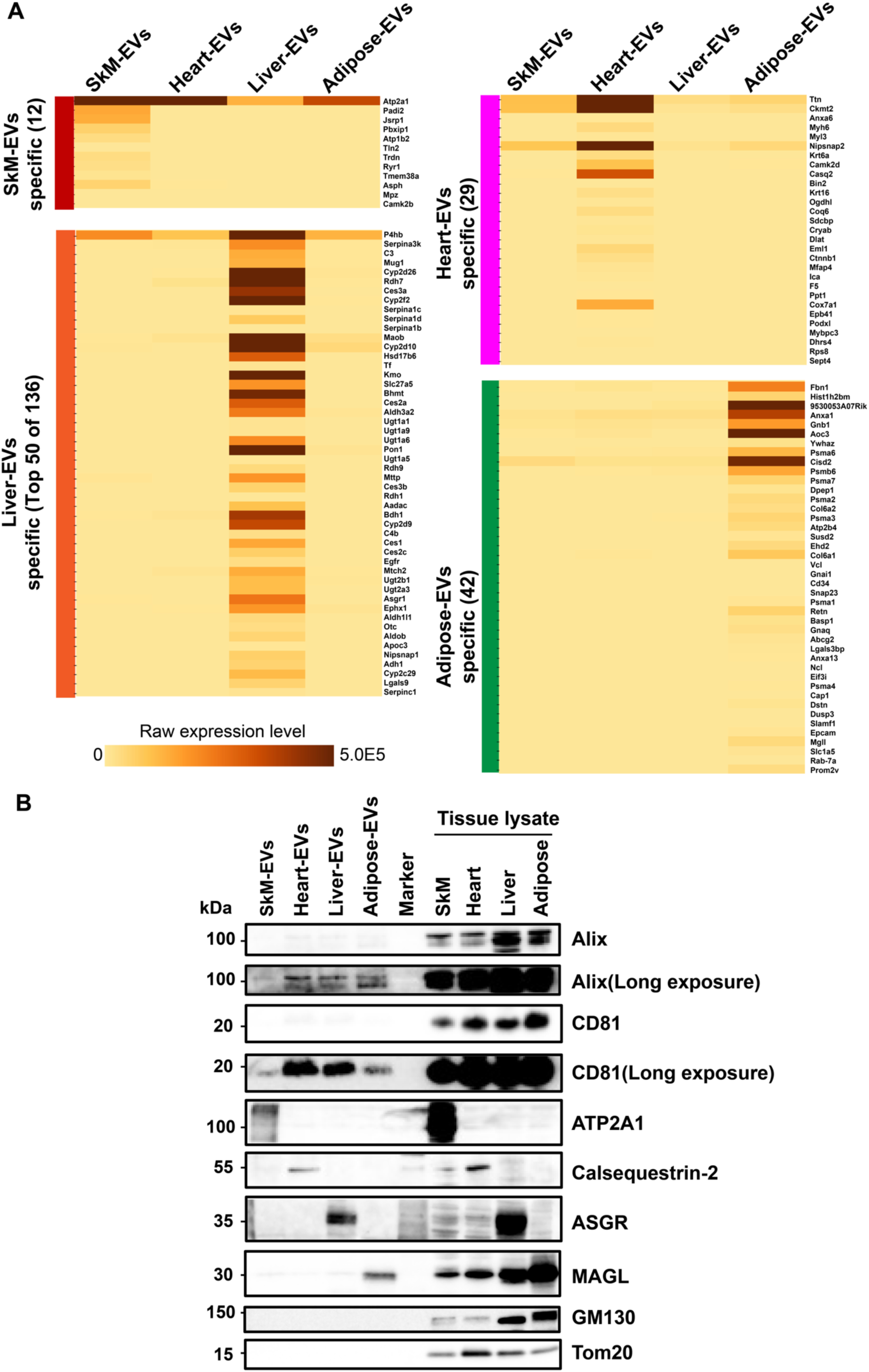
Identification of specific marker proteins of tissue EVs. (A) Heatmap showing tissue-specific enrichment of EV proteins. Tissue-specific EV proteins of the four tissues are listed. Proteins contained more than 10-fold higher in EVs from one tissue than those from the other three tissues were extracted as potential tissue-specific EV proteins. (B) Expression of classical EV marker proteins (Alix and CD81), tissue-specific EV proteins (ATP2A1, CASQ2, ASGR1, and MAGL), and non-EV proteins (GM130 and Tom20) in tissue EVs and original tissues. Tissue EVs (2 μg protein) isolated by the PS-affinity method and tissue homogenates (5 μg protein) were analyzed by immunoblotting.

To validate the bioinformatics-based results, we performed immunoblot analysis on tissue EVs from the four tissues. Tissue EVs from the four tissues contained Alix and CD81, typical EV marker proteins, whereas they were devoid of GM130 and Tom20, non-EV proteins (**Figure 4B**). Among the tissue-specific EV proteins we selected, ATP2A1, calsequestrin-2 (CASQ2), asialoglycoprotein receptor 1 (ASGR1), and monoglyceride lipase (MAGL, also known as Mgll) were detected only in SkM-EVs, heart-EVs, liver-EVs, and adipose-EVs, respectively (**Figure 4B**). We examined tissue specificities of these marker proteins by immunoblot and qRT-PCR and confirmed the tissue-specific expression of these proteins except MAGL (**Figure S5**). Collectively, the results suggest that ATP2A1, CASQ2, ASGR1, and MAGL can serve as markers for SkM-EVs, heart-EVs, liver-EVs, and adipose-EVs, respectively.

### Comparative proteome analysis of plasma and tissue EVs

Although it is hypothesized that circulating EVs are derived from virtually all tissues, how much each tissue contributes to circulating EVs is largely unknown. Finally, we sought to investigate the tissue origins of circulating EVs. To this end, we isolated plasma EVs from mice using the PS-affinity method (**Figure 5A**). The size of plasma EVs was 63.1 ± 20.6 nm, ranging from 30−120 nm (**Figure 5B**). Proteomic analysis identified a total of 464 proteins in plasma EVs (**Supplementary Table 2**). GO analysis showed the enrichment of mitochondrial-related proteins in plasma EVs (**Figure S6**), suggesting that mitochondria-derived vesicles are actively released to the bloodstream. We next performed principal component analysis (PCA) using proteomes of EVs from plasma, skeletal muscle, heart, liver, and adipose tissues. The results showed that plasma EVs are distinguishable from EVs of the four tissues, suggesting that plasma EVs are derived from various tissues and cell types (**Figure 5C**). In addition, liver-EVs also constituted a distinct group. On the other hand, SkM-EVs, heart-EVs, and adipose-EVs were categorized into the same group, indicating that proteomic profiles of EVs from these three tissues exhibit high similarity.

**Figure 5.**
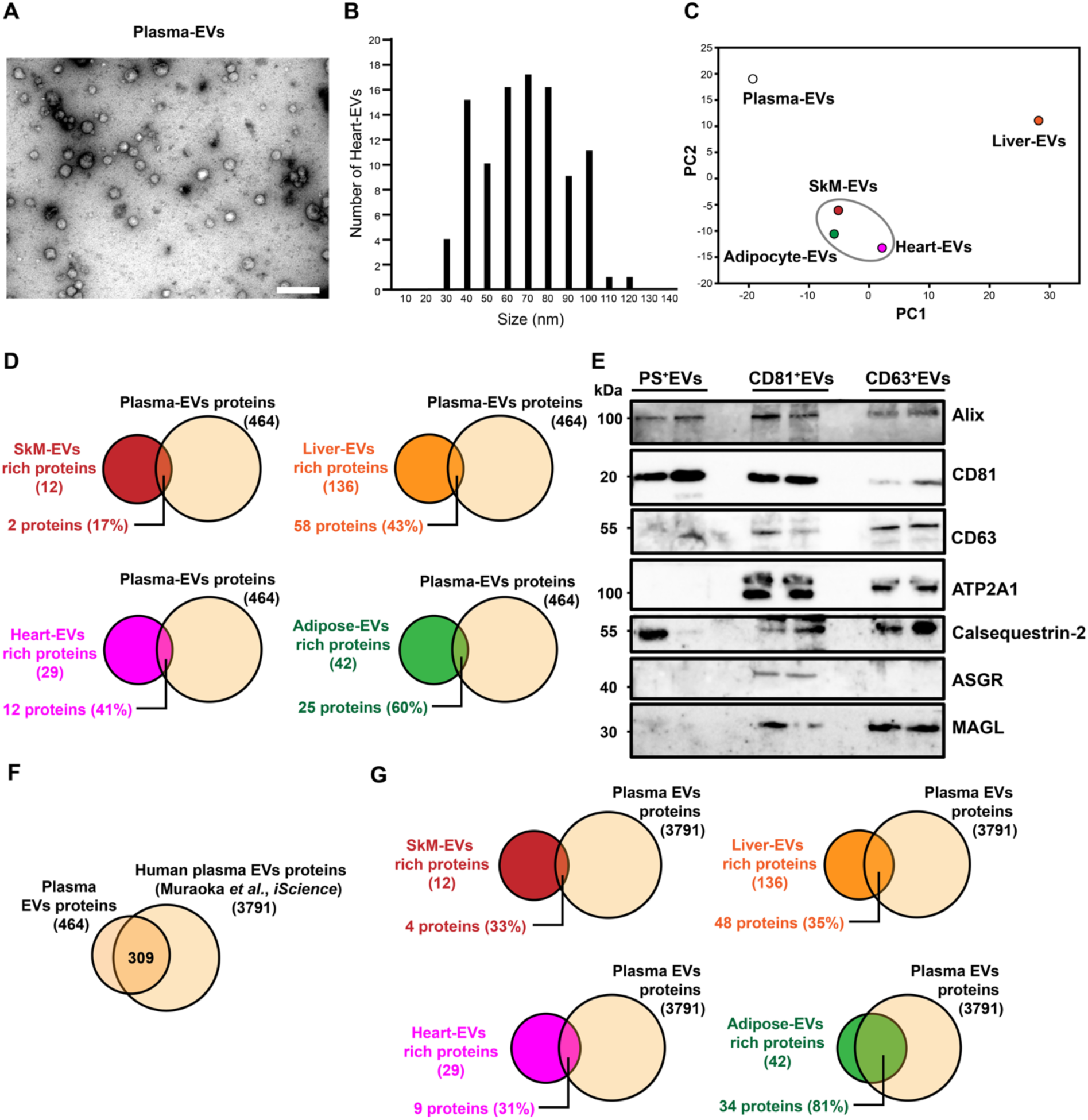
Comprehensive comparison between tissue and plasma EV proteomes. (A) TEM image of plasma EVs. Plasma EVs were isolated by PS-affinity beads as described in Materials and methods. Scale bar, 500 nm. (B) Size distribution of plasma EVs. TEM images of plasma EVs were used to analyze the size of EVs (n = 100). (C) Principal component analysis (PCA) of the proteomes of plasma, SkM, heart, liver, and adipose EVs. (D) Presence of tissue EV marker proteins in plasma EVs. Venn diagrams show the presence of tissue EV marker protein (identified in Figure 4A) and mouse plasma EV proteins. The number and percentage of overlapping proteins between tissue and plasma EVs are shown in each Venn diagram. (E) Expression of typical EV markers (Alix and CD81) and tissue-specific EV proteins (ATP2A1, CASQ2, ASGR1, MAGL) in plasma EVs. PS^+^ EVs (*left*), CD81^+^ EVs (*center*), and CD63^+^ EVs (*right*) were isolated from plasma (2 mice each) using PS-affinity beads, CD81-affinity beads, and CD63-affinity beads, respectively. Plasma EVs (2 μg protein/lane) were subjected to immunoblot analysis. (F) Venn diagram showing similarity of mouse (this study) and human (Muraoka *et al.* [17]) plasma EV proteome. Both mouse and human plasma EVs were isolated by the same PS-affinity beads. (G) Presence of tissue EV marker proteins in human plasma EVs. Venn diagrams show the presence of tissue-specific EV marker proteins of each tissue (identified in Figure 4A) in human plasma EVs [17]. The number and percentage of overlapping proteins between tissue and plasma EVs are shown in each Venn diagram.

We next asked whether the tissue-specific EV marker proteins we identified are present in plasma EVs. In 464 plasma EV proteins, two SkM-EV proteins, 12 heart-EV proteins, 58 liver-EV proteins, and 25 adipose-EV proteins were detected, which account for 17%, 41%, 43%, and 60% of tissue-EV markers, respectively (**Figure 5D**). These results suggest that tissue EVs reach the bloodstream while the accessibility to circulation varies among the four tissues. To further validate these results, we isolated mouse plasma EVs using three different methods and examined the presence or absence of tissue EV markers (**Figure 5E**); PS^+^ EVs, CD81^+^ EVs, and CD63^+^ EVs were collected by the PS-affinity, CD81-affinity, and CD63-affinity beads, respectively. The three types of plasma EVs contained the classical EV marker protein Alix, confirming the isolation of EVs by the three methods. As expected, the isolation of plasma EVs with CD81-affinity beads and CD63-affinity beads yielded CD81-rich and CD63-rich EVs. PS^+^ EVs were relatively rich in CD81 compared to CD63. We next examined whether tissue EV marker proteins exist in the three types of plasma EVs. The results showed that the contents of tissue EV marker proteins vary among PS^+^ EVs, CD81^+^ EVs, and CD63^+^ EVs; the SkM-EV marker protein ATP2A1 and the adipose-EV marker protein MAGL were detected in CD81^+^ EVs and CD63^+^ EVs, while ASGR1, a liver-EV marker protein, was detected only in CD81^+^ EVs. The heart-EV marker protein CASQ2 was present in all three types of plasma EVs. Collectively, our findings suggest that plasma EVs are a highly heterogeneous population of different origins. In addition, it is suggested that protein cargoes of EVs, even from the same tissue of the origin, are significantly different.

Finally, we sought to examine whether the tissue EV marker proteins identified above are applicable to assessing sources of human plasma EVs. We utilized a proteome dataset of human plasma EVs recently reported [17] because human plasma EVs were isolated by the same PS-affinity method as ours in the report. We first compared proteomes between mouse (this study) and human (Muraoka *et al.* [17]) plasma EVs. Sixty-seven % of mouse plasma PS^+^ EV proteins, which represent 309 out of 464 proteins, were found in human plasma PS^+^ EVs (**Figure 5F**), indicating similar protein profiles of plasma EVs in mice and humans. We next asked whether human plasma EVs contain our tissue EV marker proteins. The results show that four SkM-EV, nine heart-EV, 48 liver-EV, and 34 adipose-EV marker proteins are present in human plasma EVs, which account for 33%, 31%, 35%, and 81% of tissue-specific marker proteins, respectively (**Figure 5G**). These results suggest that EVs from adipose tissues enter circulation more readily than skeletal muscle, heart, and liver not only in mice but also in humans, and that the accessibility of tissue EVs to the bloodstream differs among tissues.

## Discussion

Although it is proposed that EVs mediate interorgan communications through systemic circulation, little is known about whether each tissue equally contributes to circulating EVs. In addition, whether potential tissue EV markers identified by cell-based studies are suitable for monitoring the transport of tissue EVs in the body is controversial [39, 40]. We hypothesized that EVs of a certain tissue are highly concentrated in the interstitium of the tissue. Furthermore, we expected that proteomic profiling of EVs of each tissue facilitates predicting the origin of circulating EVs.

Because developing a solid method for isolating EVs from different tissues should be an important technical advance in determining proteome of tissue EVs and investigating EVs *in vivo*, we first developed methods to collect EVs from different tissues, including skeletal muscle, heart, liver, and adipose tissues. Although several groups have independently established protocols for isolating EVs from cancer tissues [22], adipose tissues [23], heart [34], and brain [41, 42], these methods have not been validated in different tissues. In this work, we refined previous methods and developed the UC-digestion and PS-affinity methods for efficiently collecting EVs from different tissues. The two methods have both advantages and disadvantages. The UC-digestion recovers the bulk of EVs from tissues, being suitable for studying their overall characteristics. This method has several limitations, including co-precipitation of small amounts of undigested ECM and the aggregation of EVs. On the other hand, the PS-affinity method can isolate intact EVs without the inclusion of ECM and the aggregation of EVs. However, the PS-affinity method is unable to collect PS-negative EVs [43]. Various types of cells exist in a tissue, and each cell type and presumably even the same cell type release EVs with different cargoes [44–47].

Therefore, further modifications such as the combination with size exclusion chromatography [48], tetraspanin-affinity beads [49], or antibodies to cell type-specific proteins will help isolate more specific EVs of interest.

The lack of reliable biomarkers for tissue EVs has limited our understanding of *in vivo* functions of tissue EVs [39]. In this study, we determined proteomic profiles of EVs of skeletal muscle, heart, liver, and adipose tissues, and identified ATP2A1, CASQ2, ASGR1, and MAGL as marker proteins for skeletal muscle, heart, liver, and adipose EVs, respectively. Previous studies support our findings; ATP2A1 and ASGR1 were identified as skeletal muscle and liver EV markers, respectively [19, 50, 51]. On the other hand, this study is the first to propose that Calsequestrin 2 and MAGL serve as heart and adipose EVs, respectively. In addition, because ATP2A1 and ASGR1 are transmembrane proteins, it is worth noting that antibodies to an extravesicular domain of these proteins may enable us to specifically isolate skeletal muscle and liver EVs from biological fluids.

Based on our proteomic profiling of tissue and plasma EVs, we found that mitochondria-related proteins were unexpectedly enriched not only in tissue EVs but also plasma EVs. Recent studies showed that small EVs contain mitochondrial proteins as cargoes in the heart [34], brain [35, 41], and adipose tissues [52, 53]. Further investigation is needed to clarify whether the mitochondrial proteins are secreted to eliminate the damaged mitochondria as EVs, particular mitochondrial proteins are selectively incorporated into EVs, and certain mitochondrial proteins are actively transferred to recipient cells [33]. However, consistent with our proteomics and immunoblot data (**Supplementary Table 2**, **Figure 4B**), Liang *et al.* showed that the mitochondrial outer membrane protein Tom20 was undetectable in small EVs isolated from heart tissue. These results suggest the selective incorporation of mitochondrial proteins into EVs.

The PCA of our proteomic data on tissue and plasma EVs revealed the clustering of SkM-EVs, heart-EVs, and adipose-EVs (**Figure 5C**). Given that skeletal muscle, heart, and adipose tissues originate from the mesoderm while the liver is from the endoderm [54], the tissue lineage may reflect protein components of tissue EVs. Indeed, previous studies suggest that the similarities of tissue protein profiles often coincide with the putative developmental origin of cell types [55]. The current analysis also shows that the plasma EV proteome is distinct from all tissue EVs tested, suggesting that circulating EVs are of various different origins or from other tissues rather than the four tissues we examined.

We sought to apply our proteomic data on tissue and plasma EVs to predict the accessibility of EVs from a tissue to circulation by comprehensive comparison. The current analysis suggests that, among the four tissues, adipose-EVs readily enter the bloodstream whereas skeletal muscle exhibits the smallest contribution to the plasma EVs. Our prediction is supported by previous findings; it has been shown that skeletal muscle is not the major source of circulating EVs despite the largest organ in the body [19, 56, 57]. On the other hand, studies with adipocyte-specific Dicer knockout mice show that adipose tissues are the major source of circulating miRNA, suggesting that the predominant origin of circulating EVs is adipose tissues [58]. Moreover, adipose tissue-derived EVs have been detected in human and mouse plasma [59, 60].

In summary, we determined proteomic profiling and tissue-specific marker proteins of tissue EVs by developing protocols to isolate EVs from different tissues. Multiple bioinformatic comparisons suggest that the accessibility of EVs to the bloodstream is highly dependent on the tissue of origin. It is expected that our tissue EV proteomes and marker proteins serve as valuable resources for investigating EVs *in vivo*.

## Materials and Methods

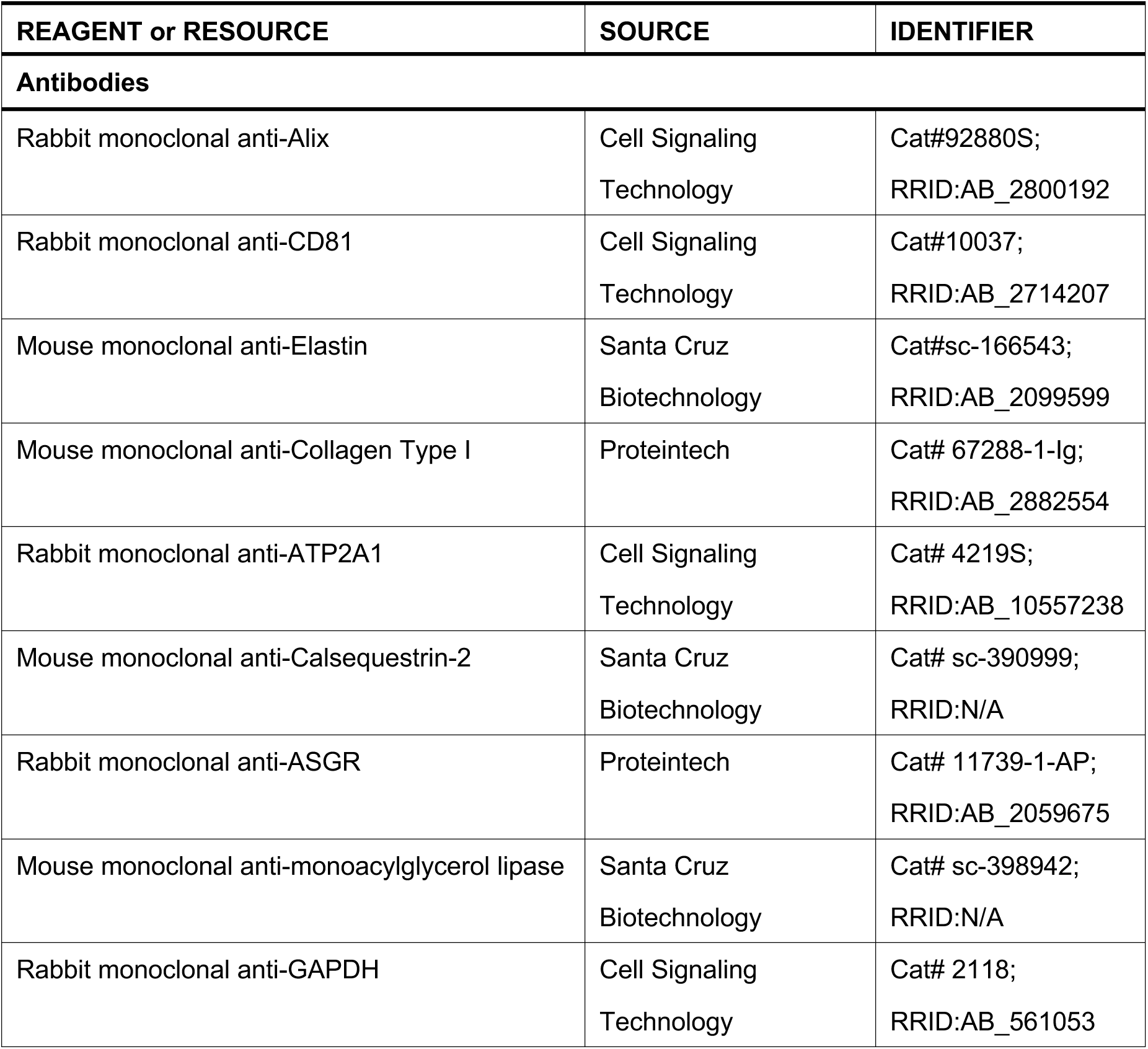
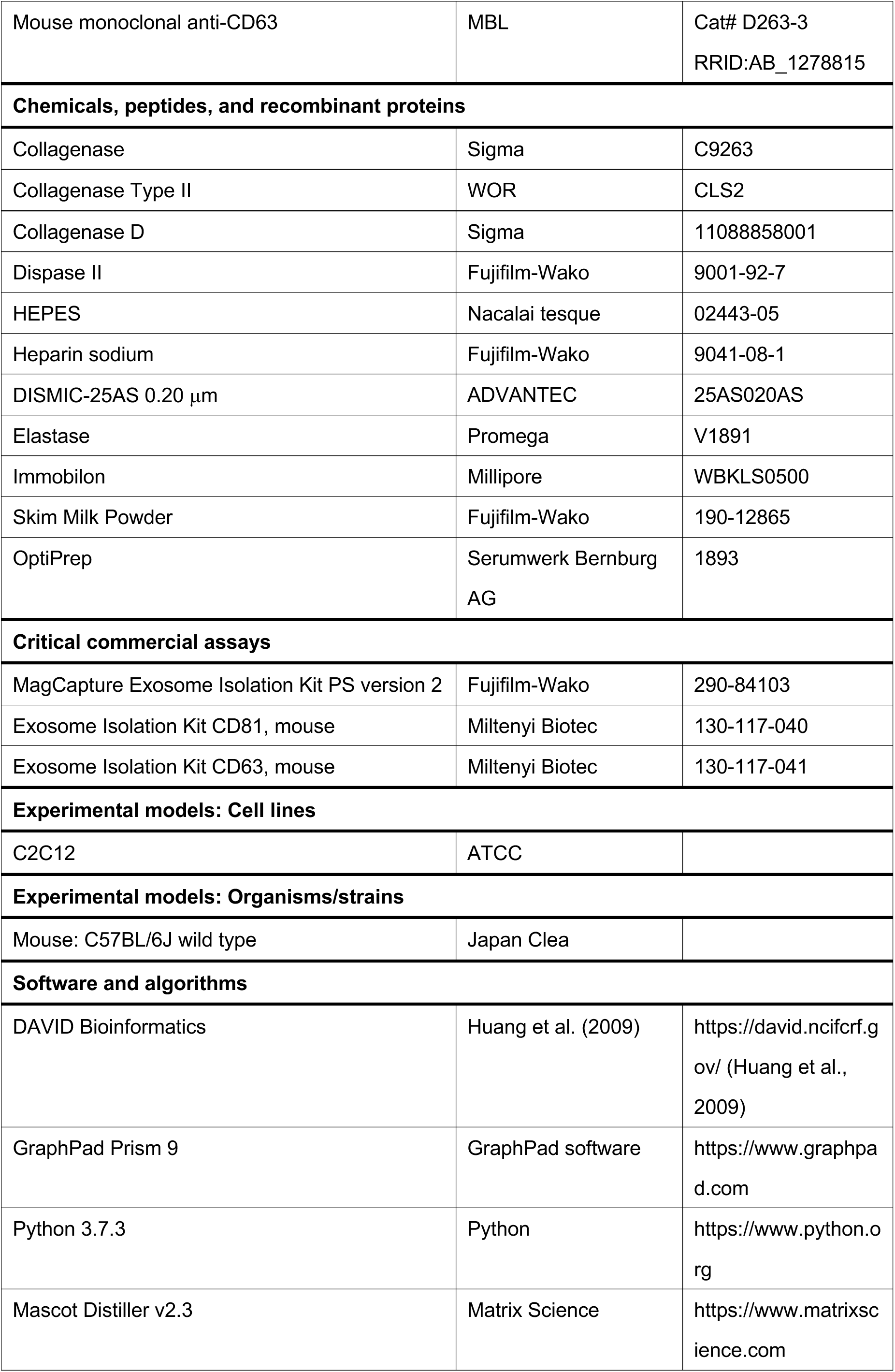
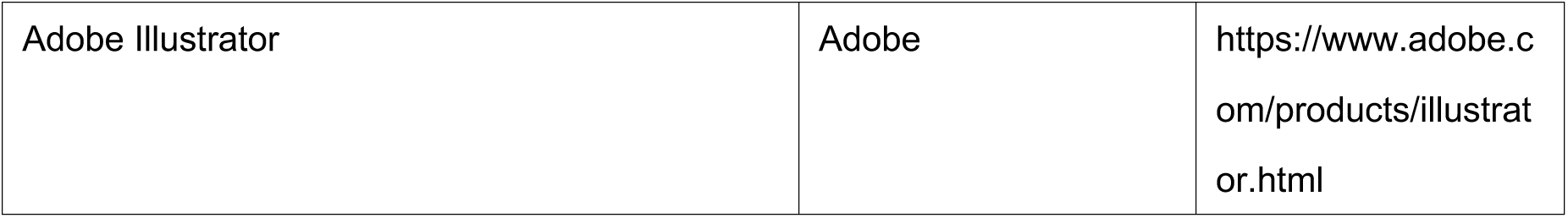
Key Resource Table

Key Resource Table
| REAGENT or RESOURCE | SOURCE | IDENTIFIER |
| --- | --- | --- |
| **Antibodies** |  |  |
| Rabbit monoclonal anti-Alix | Cell Signaling Technology | Cat#92880S;<br>RRID:AB_2800192 |
| Rabbit monoclonal anti-CD81 | Cell Signaling Technology | Cat#10037;<br>RRID:AB_2714207 |
| Mouse monoclonal anti-Elastin | Santa Cruz Biotechnology | Cat#sc-166543;<br>RRID:AB_2099599 |
| Mouse monoclonal anti-Collagen Type I | Proteintech | Cat# 67288-1-Ig;<br>RRID:AB_2882554 |
| Rabbit monoclonal anti-ATP2A1 | Cell Signaling Technology | Cat# 4219S;<br>RRID:AB_10557238 |
| Mouse monoclonal anti-Calsequestrin-2 | Santa Cruz Biotechnology | Cat# sc-390999;<br>RRID:N/A |
| Rabbit monoclonal anti-ASGR | Proteintech | Cat# 11739-1-AP;<br>RRID:AB_2059675 |
| Mouse monoclonal anti-monoacylglycerol lipase | Santa Cruz Biotechnology | Cat# sc-398942;<br>RRID:N/A |
| Rabbit monoclonal anti-GAPDH | Cell Signaling Technology | Cat# 2118;<br>RRID:AB_561053 |
| Mouse monoclonal anti-CD63 | MBL | Cat# D263-3<br>RRID:AB_1278815 |
| **Chemicals, peptides, and recombinant proteins** |  |  |
| Collagenase | Sigma | C9263 |
| Collagenase Type II | WOR | CLS2 |
| Collagenase D | Sigma | 11088858001 |
| Dispase II | Fujifilm-Wako | 9001-92-7 |
| HEPES | Nacalai tesque | 02443-05 |
| Heparin sodium | Fujifilm-Wako | 9041-08-1 |
| DISMIC-25AS 0.20 μm | ADVANTEC | 25AS020AS |
| Elastase | Promega | V1891 |
| Immobilon | Millipore | WBKLS0500 |
| Skim Milk Powder | Fujifilm-Wako | 190-12865 |
| OptiPrep | Serumwerk Bernburg<br>AG | 1893 |
| **Critical commercial assays** |  |  |
| MagCapture Exosome Isolation Kit PS version 2 | Fujifilm-Wako | 290-84103 |
| Exosome Isolation Kit CD81, mouse | Miltenyi Biotec | 130-117-040 |
| Exosome Isolation Kit CD63, mouse | Miltenyi Biotec | 130-117-041 |
| **Experimental models: Cell lines** |  |  |
| C2C12 | ATCC |  |
| **Experimental models: Organisms/strains** |  |  |
| Mouse: C57BL/6J wild type | Japan Clea |  |
| **Software and algorithms** |  |  |
| DAVID Bioinformatics | Huang et al. (2009) | https://david.ncifcrf.gov/ (Huang et al., 2009) |
| GraphPad Prism 9 | GraphPad software | https://www.graphpad.com |
| Python 3.7.3 | Python | https://www.python.org |
| Mascot Distiller v2.3 | Matrix Science | https://www.matrixscience.com |
| Adobe Illustrator | Adobe | https://www.adobe.com/products/illustrator.html |

### Mice

All protocols for animal procedures were approved by the Animal Care and Use Committee of the University of Tokyo (Approval number P16-266), which are based on the Law for the Humane Treatment and Management of Animals (Law No. 105, 1 October 1973, as amended on 1 June 2020). Male C57BL/6J mice at 8 weeks old were obtained from Japan Clea. Mice were housed in a 12 h-light/12 h-dark schedule at 23 ± 2°C and 55 ± 10% humidity and fed *ad libitum* with a standard chow diet (Labo MR Stock, Nosan Corporation) and water. Mice at 9 to 10 weeks old were perfused through the heart at the left ventricle with PBS at a rate of 1 ml/min for 2 min to remove blood from the tissue under anesthesia with isoflurane. Afterward, skeletal muscle (tibialis anterior, gastrocnemius, soleus, quadriceps), liver, heart, and epididymal white adipose tissues were then harvested.

### Isolation of tissue EVs by ultracentrifugation

Tissue EVs were isolated based on methods recently reported [22, 23] with modifications. Approximately 600 mg of skeletal muscle tissues (mixed of tibialis anterior, gastrocnemius, soleus, and quadriceps) from a hind limb, heart, liver, and epididymal white adipose tissue were each dissected and digested with collagenase (10 mg/mL, Sigma) and dispase II (10,000 PU/mL, Fujifilm-Wako) for 1 h at 37°C in HEPES buffer (100 mM HEPES pH 7.4, 2.5 mM CaCl_2_). To avoid disruption of cells, tissues were minced gently. Afterward, one volume of PBS containing 2 mM EDTA was added to the sample, and the sample was passed through a 100 μm cell strainer (Corning). Samples were then centrifuged at 600 g for 5 min at 4°C, 2,000 g for 10 min, and 10,000 g for 30 min. The supernatant was filtrated with a 0.20 μm filter (Advantec) to obtain small EVs. EVs were pelleted by ultracentrifugation at 100,000 g for 70 min at 4°C (Beckman-Coulter). The EVs pellet was washed once with PBS (2 mL/tube) and EVs were pelleted by ultracentrifugation at 100,000 g for 70 min at 4°C again. The collecting pellet was resuspended in 150 μL of PBS. For Optiprep density-gradient centrifugation, 1 mL of different concentrations of Optiprep (47, 37, 28, and 18% (w/v) diluted with 20 mM HEPES, pH 7.4) were sequentially layered, and EVs isolated by ultracentrifugation were overlaid on the top. After ultracentrifugation for 16 h at 4°C at 200,000 g, eight 500-μL fractions were collected from the top. Fractions were diluted with 2 mL PBS, and the volume of a sample was reduced to 150 μL by ultrafiltration (Amicon Ultra 0.5 ML – 100kDa, Merck) for exchanging buffer.

### Isolation of SkM-EVs by UC-digestion method

Tissue EVs were collected from skeletal muscle tissues by ultracentrifugation as described above. 50 μL of EV samples were resuspended by 1 mL of Tris-HCl buffer (50 mM Tris-HCl pH 9.0, 2.5 mM CaCl_2_), and subjected to additional digestion with collagenase (10 mg/mL, Sigma) and Elastase (5 mg/mL, Promega) for 1 or 3 h at 37°C. After the digestion, EVs were pelleted by ultracentrifugation at 100,000 g for 70 min at 4°C twice. The resulting pellet was resuspended in 150 μL of PBS. EV protein contents were determined by the Micro BCA Protein Assay Kit (Thermo Fisher). EVs were stored at -80°C until use.

### Isolation of plasma and tissue EVs using affinity beads

Plasma EVs were isolated by MagCapture Exosome Isolation Kit PS (Fujifilm-Wako), Exosome Isolation Kit CD63, mouse (Miltenyi Biotec), or Exosome Isolation Kit CD81, mouse (Miltenyi Biotec) according to the manufacturer’s procedures. Plasma was collected by adding heparin sodium solution (Fujifilm-Wako) to the blood (Final concentration: 5 U/mL). Plasma (300 μL) was mixed with PBS (600 μL) and spun at 10,000 g for 30 min at 4°C. After filtration of the supernatant with a 0.20 μm filter, plasma sample was subjected to the isolation of EVs using MagCapture Exosome Isolation Kit PS, Exosome Isolation Kit CD63, or Exosome Isolation Kit CD81 according to the manufacturer’s procedures. For tissue EVs isolation, the 10,000-g supernatant was prepared as described above. After filtration with a 0.20 μm filter and concentration with Amicon Ultra-15 (Merck), EVs were isolated using MagCapture Exosome Isolation Kit PS (Fujifilm-Wako). In brief, plasma or tissue supernatant containing small EVs (∼1 mL) was mixed with 0.6 mg of streptavidin magnetic beads bound to 1 μg of biotinylated mouse Tim 4-Fc and incubated for 16–18 h in the presence of 2 mM CaCl_2_ at 4°C with rotation. After washing beads three times with 1 mL of washing buffer (20 mM Tris-HCl pH 7.4, 150 mM NaCl, 2 mM CaCl_2_, 0.0005% Tween 20), EVs were eluted twice with 50 μl of elution buffer (20 mM Tris-HCl pH 7.4, 150 mM NaCl, 2 mM EDTA). Plasma and tissue EVs were eluted with an elution volume of 200 μL per 300 μL plasma and 200 μL per tissue sample, respectively.

### Transmission electron microscopy

Specimens for transmission electron microscopy were prepared at room temperature. An aliquot of EV sample was pipetted onto a copper grid with carbon support film and incubated for 10 min. After removing the excess liquid, a grid was briefly placed on 10 μL of 2% uranyl acetate (w/v, Merck). Images were acquired under a JEM-1010 electron microscope (JEOL) operated at 100 kV with a Keen view CCD camera (Olympus Soft Imaging Solution). The size of EVs was measured using Image J software.

### Immunoblotting

Tissue homogenates were prepared in radioimmunoprecipitation assay buffer (50 mM Tris-HCl pH 7.4, 150 mM NaCl, 1 mM EDTA, 1% Nonidet P-40, and 0.25% sodium deoxycholate) supplemented with a protease inhibitor mixture (Nacalai tesque) and phosphatase inhibitor mixture (Sigma). Protein concentration was determined by BCA Protein Assay (Thermo Fisher). Tissue homogenate or EVs were mixed with Laemmli buffer and heated at 70°C for 10 min. Aliquots were subjected to SDS-PAGE and immunoblotting according to a standard protocol. The expression of protein of interest was analyzed using Image J software or Evolution-Capt software (Vilber Lourmat).

### mRNA expression analysis

Total RNA was isolated using ISOGEN (NIPPON GENE) according to the manufacturer’s instructions. The high-capacity cDNA reverse transcription kit (Applied Biosystems) was used to synthesize cDNA from total RNA. Quantitative real-time PCR (qRT-PCR) analyses were performed using an Applied Biosystems StepOnePlus. mRNA levels were normalized to 18S ribosomal RNA levels as described [19]. The primers used for qPCR are listed in **Supplementary Table 5**.

### Proteomic analysis of EVs

Proteomic analysis was conducted as described [19]. EVs were solubilized in 50 mM Tris-HCl pH 9.0 containing 5% sodium deoxycholate, reduced with 10 mM dithiothreitol for 60 min at 37°C, and alkylated with 55 mM iodoacetamide for 30 min in the dark at 25°C. The reduced and alkylated samples were diluted 10-fold with 50 mM Tris-HCl pH 9.0 and digested with trypsin at 37°C for 16 h (trypsin-to-protein ratio of 1:20 (w/w)). An equal volume of ethyl acetate was added to each sample solution, and the mixtures were acidified with the final concentration of 0.5% trifluoroacetic acid. The mixtures were shaken for 1 min and centrifuged at 15,700 g for 2 min. The aqueous phase was collected and desalted with C18-StageTips. LC-MS/MS analysis was performed using an UltiMate 3000 Nano LC system (Thermo Fisher) coupled to Orbitrap Fusion Lumos hybrid quadrupole-Orbitrap mass spectrometer (Thermo Fisher) with a nano-electrospray ionization source. The sample was injected by an autosampler and enriched on a C18 reverse-phase trap column (100 μm × 5 mm length, Thermo Fisher) at a flow rate of 4 μL/min. The sample was subsequently separated by a C18 reverse-phase column (75 μm × 150 mm length, Nikkyo Technos) at a flow rate of 300 nL/min with a linear gradient from 2 to 35% mobile phase B (95% acetonitrile and 0.1% formic acid). The peptides were ionized using nano-electrospray ionization in positive ion mode.

### MS data analysis

The raw data were analyzed by Mascot Distiller v2.3 (Matrix Science), and peak lists were created based on the recorded fragmentation spectra. Peptides and proteins were identified by Mascot v2.3 (Matrix Science) using the UniProt database with a precursor mass tolerance of 10 ppm, a fragment ion mass tolerance of 0.01 Da, and strict trypsin specificity allowing for up to 1 missed cleavage. The carbamidomethylation of cysteine and the oxidation of methionine were allowed as variable modifications.

### Statistical analysis

GraphPad Prism 9 was used to analyze and plot all data. One-way ANOVA with Tukey-Kramer post-hoc test was conducted to determine the statistical significance of results. P values less than 0.05 were considered statistically significant.

## Acknowledgments

We thank Drs. Satoshi Kimura and Kenji Tomita for their assistance with transmission electron microscopy.

## Supporting information

This article contains supplementary information.

## Data availability

All data are included in the article and Supplementary Information.

## Authors contributions

Y.Y. and S.W. conceived and designed the research; S.W. performed all experiments with suggestions from Y.Y., R.S., T.S., and Y.T.; S.W., Y.Y., and R.S. analyzed data; S.W. and Y.Y. wrote and edited the manuscript. All authors read and approved the manuscript.

