## Supplementary Figures for "Proteomic profiling of tissue extracellular vesicles (EVs) identifies tissue-specific EV markers and predicts the accessibility of tissue EVs to the circulation"

Supplementary information contains:

Figures S1–S6

Tables S1–S5

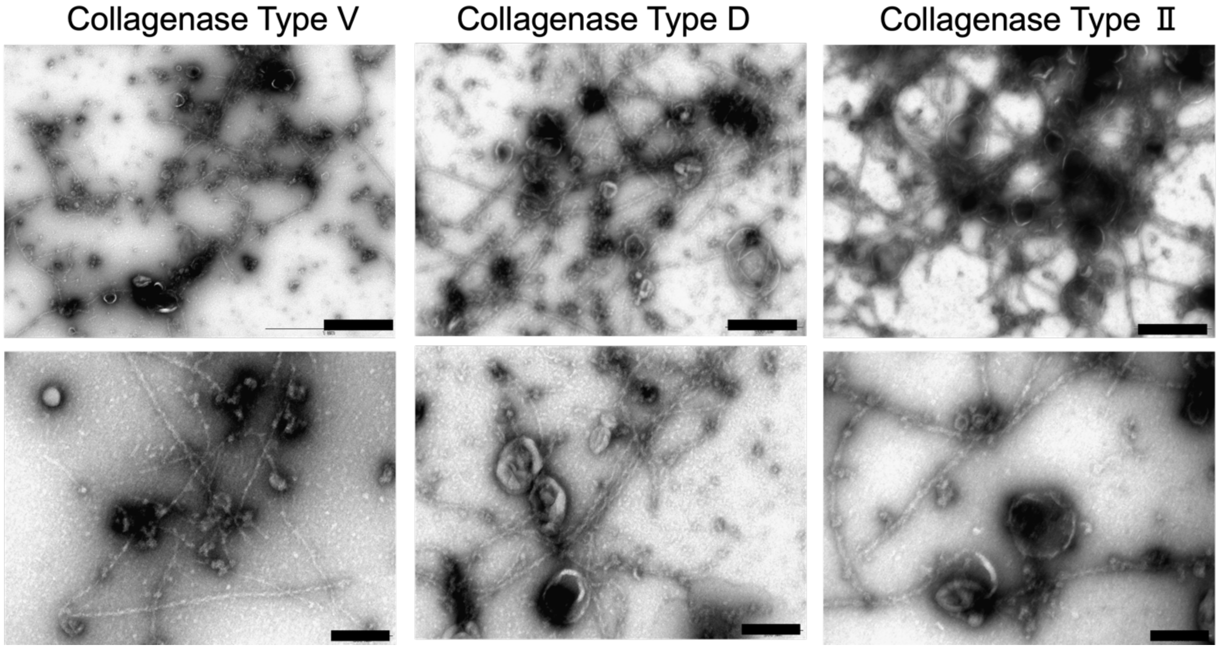


**Figure S1**

**Different proteases do not affect the digestion efficiency of ECM-like structures in SkM-EVs**

TEM images of SkM-EVs isolated using the UC method after the treatment with the indicated proteases. Scale bar; 500 nm (upper), 200 nm (lower).

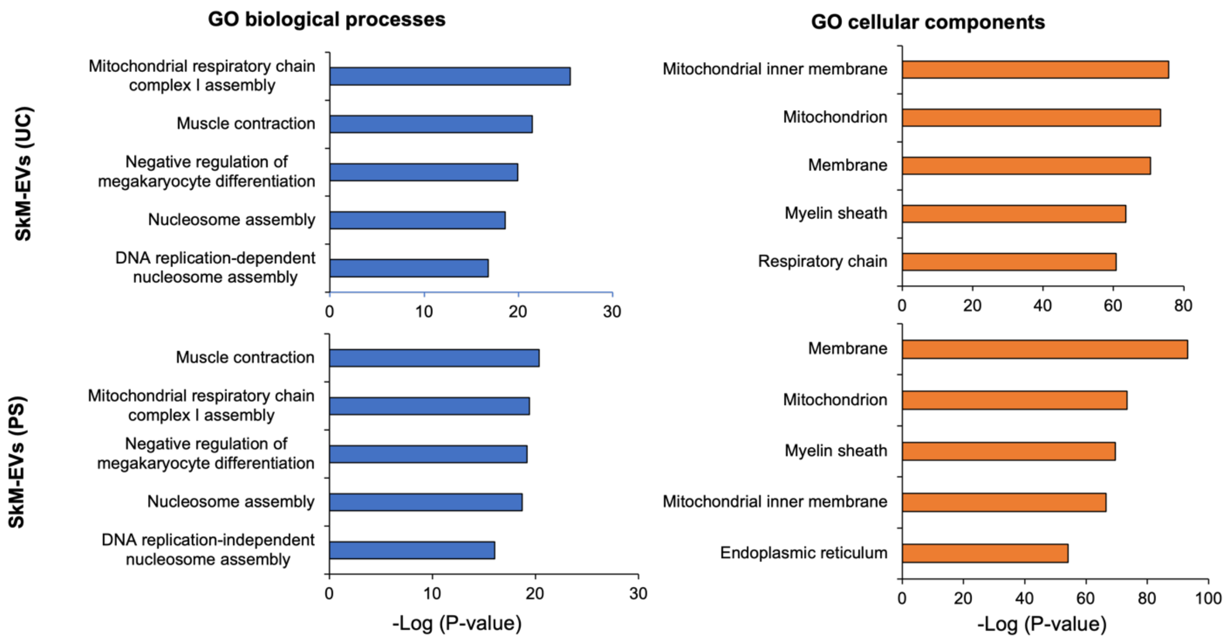


**Figure S2**

**GO analysis of SkM-EVs isolated by the UC-digestion and PS-affinity methods**

GO analyses (left, biological process; right, cellular components) were performed on SkM-EVs. SkM-EVs were isolated by the UC digestion (upper) and PS methods (lower). Proteomic data were analyzed using DAVID. The top 5 terms are listed.

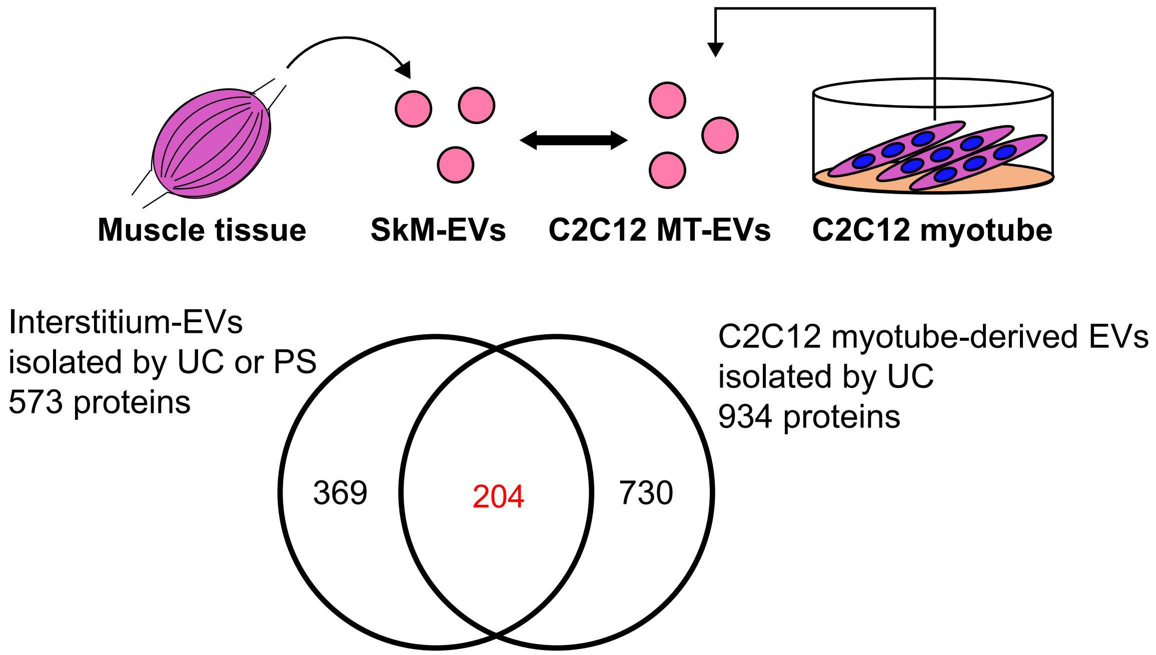


**Figure S3**

**Proteomic profiling of SkM-EVs and C2C12 myotube-derived EVs**

Venn diagram shows the distinct and overlapping EV proteins of SkM-EVs and C2C12 myotube-derived EVs. Proteomic data from SkM-EVs isolated by the UC-digestion and PS-affinity methods (this study) and from C2C12-derived EVs isolated by ultracentrifugation (reported in [1]) were used.

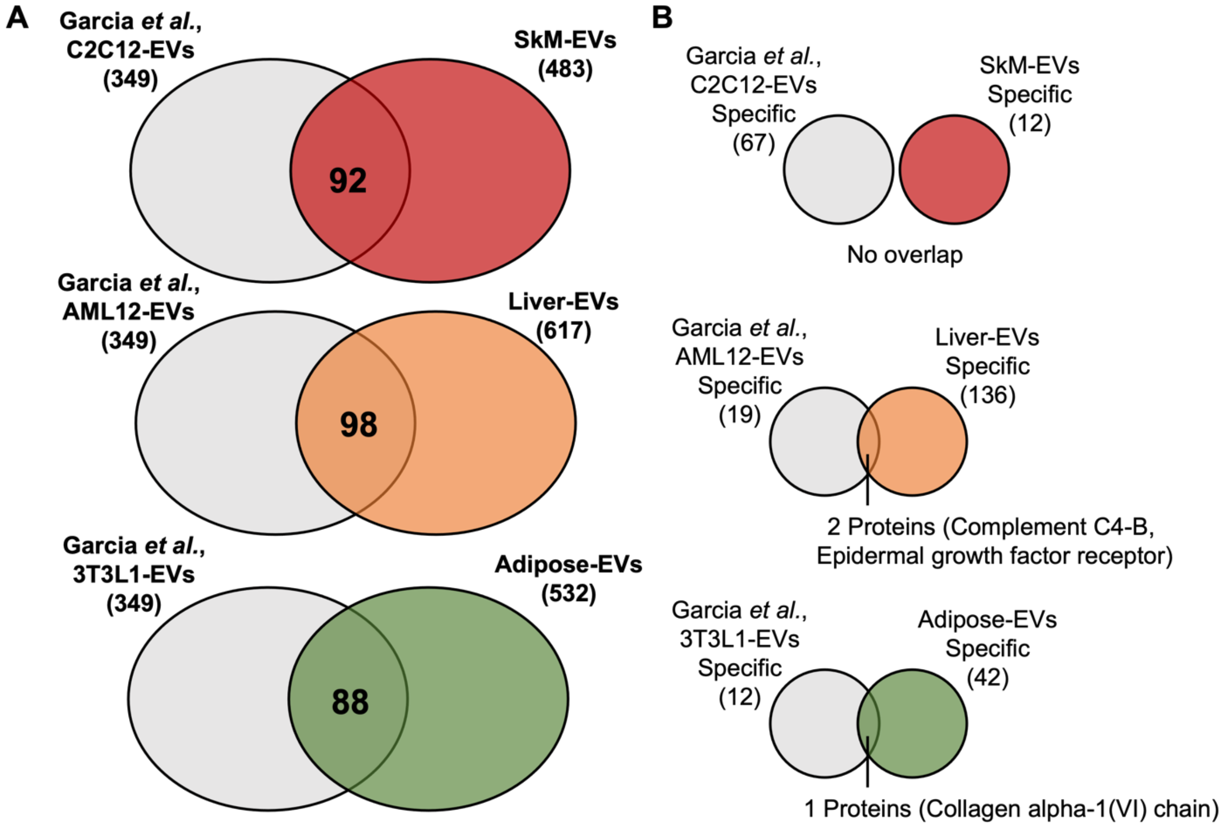


**Figure S4**

**Proteomic comparison between tissue EVs and cell line-derived EVs**

(A) Venn diagrams showing proteins overlapping between tissue EVs and EVs released from cell lines reported by Garcia *et al*. [2] (349 proteins in total).

(B) Venn diagrams showing overlap between tissue-specific EV proteins identified in this study and cell line-specific EV proteins identified by Garcia *et al*. [2].

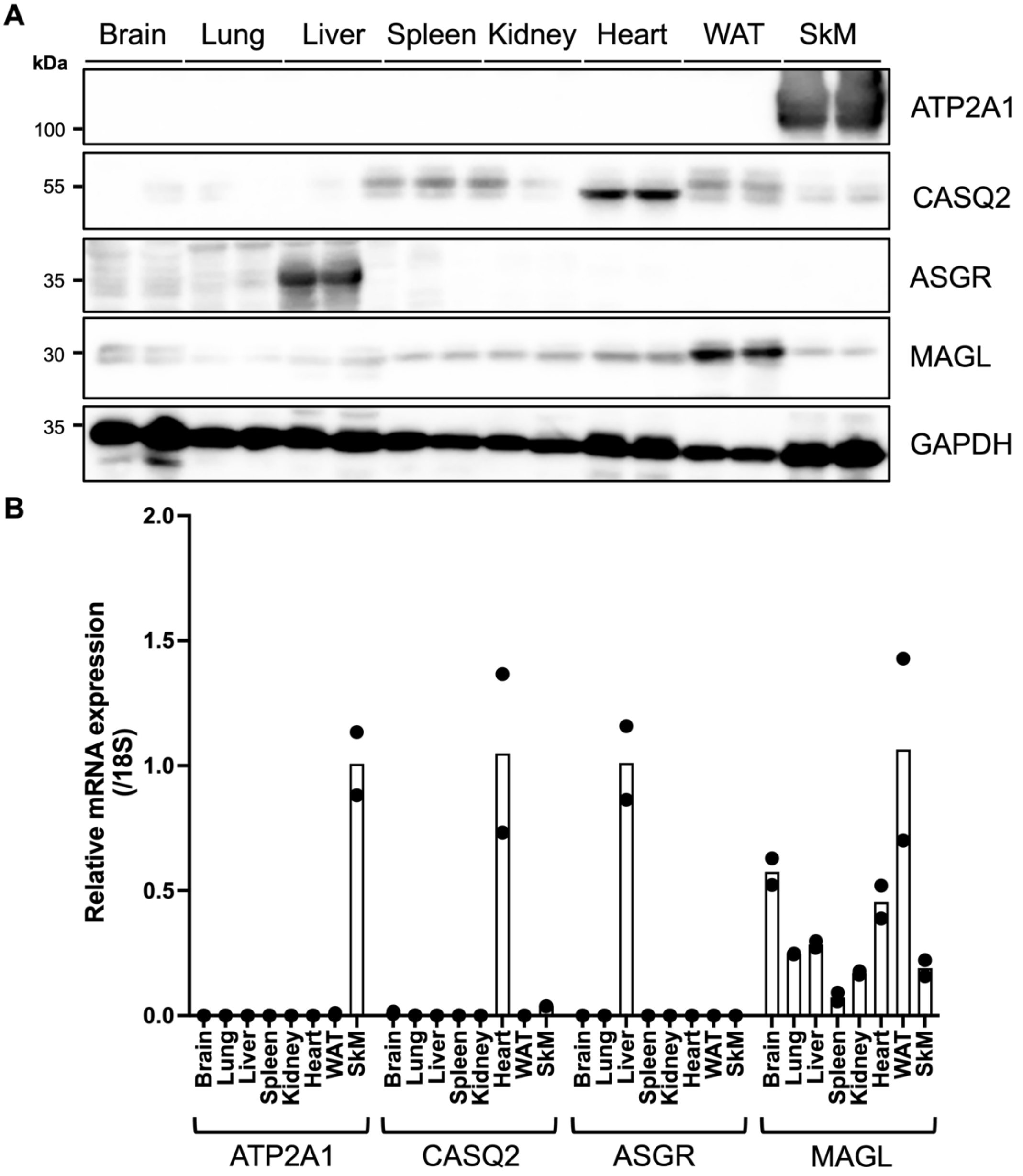


**Figure S5**

**Protein and mRNA expression of tissue-specific EV markers in mouse tissues**

(A) Protein expression. Tissue homogenates were prepared as described in Materials and Methods. Equal amounts of proteins (10 μg/lane) were subjected to immunoblot analysis using the indicated antibodies. GAPDH was detected as a loading control.

(B) mRNA levels of marker proteins. qRT-PCR was performed to determine mRNA levels of marker proteins. Results shown are means with individual data points (n=2).

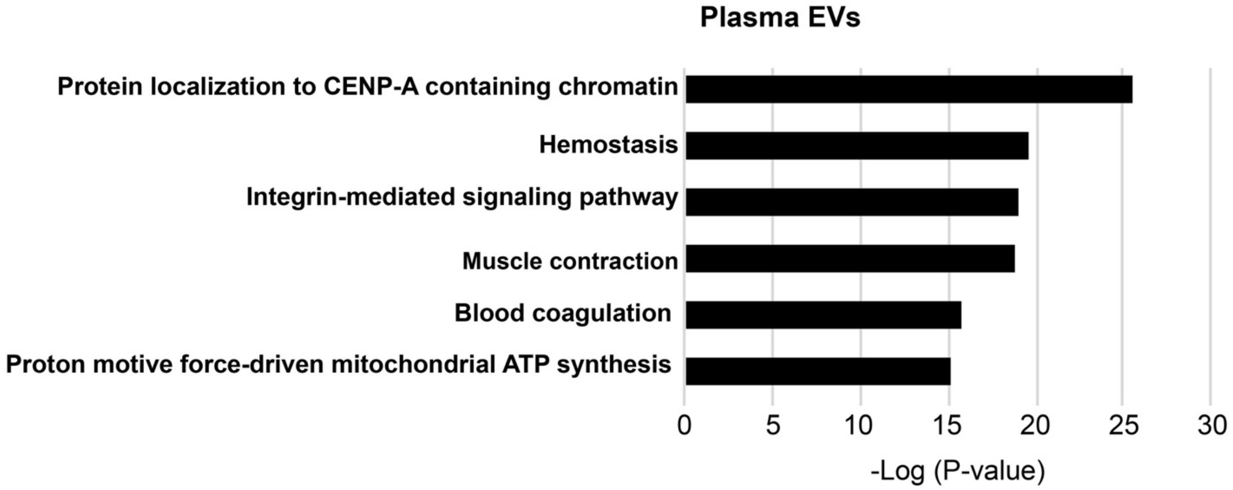


**Figure S6**

**GO analysis of plasma EVs isolated by the PS-affinity method**

GO analysis (biological process) of mouse plasma EVs. Top 6 biological process terms are listed. Proteomic data were analyzed using DAVID.
